# The Microglia Forebrain Assembloid Model Recapitulates Human Brain Development and Neuroimmune Biology

**DOI:** 10.64898/2026.04.08.717256

**Authors:** Zeinab Tashi, Kaila Gemenes, Santiago Ochoa, Mauri Spendlove, Mischa Ellison, Rose Graf, Madeline G. Andrews, Benjamin B. Bartelle

## Abstract

Microglia are innate immune cells of the CNS whose dysfunction contributes to inflammation and metabolic changes across neurodegenerative and CNS disorders. Across all stages of life, microglia are essential for immune surveillance, neural homeostasis, and synaptic pruning; however, their role in neurodevelopment is less understood. Microglia invade the brain during early neurogenesis, prior to neuronal/glial differentiation, but their potential role at this stage remains undescribed. To model neuroimmune interactions during human cortical development, we created an “assembloid” of human ESC–derived forebrain organoids combined with developmentally matched microglia during cortex formation. Functional contributions of microglia were compared to control organoids using histology and metabolomics.

## INTRODUCTION

Human induced stem cell (hiSC) derived forebrain organoids are powerful 3D tissue-like platforms for mechanistic studies of human biology. Their heterogeneous cell composition is crucial for neuron-glial interactions, but the human relevance of this biology is determined by how accurately the neural tissues recapitulate human development. Cortical expansion and organization are established during gestation as neural crest cell populations diversify and differentiate into functionally distinct neurons and glia, but the brain, as an organ, consists of other cell types that are not neural crest derived and arrive from peripheral tissues.

One cell type that invades neural tissues just as tissue patterning begins are microglia, the resident tissue immune cell of the brain. These myeloid lineage cells, similar to macrophages, originate from the yolk sac and migrate from the liver during early angiogenesis in all vertebrates (Ginhoux et al., 2010)(Wu et al., 2025). Throughout all stages of life, microglia serve immune functions, however they also serve important roles in neural tissue development, with myelination, and synapse pruning continuing into adulthood (Paolicelli et al., 2022).

Given the severe developmental phenotypes of microglia loss in human development, there are likely other roles these cells play in development. The potential to recapitulate microglia co-patterning in heterogeneous tissues makes organoid/microglia assembloid models a powerful tool for discovery. To this end primary microglia have been assembled with a forebrain organoid at the beginning of neural differentiation resulting in a tissue model that is transcriptomicaly similar to prenatal brain. While we have not yet recapitulated the yolk sac differentiation lineage of microglia in vitro, bone marrow lineage myeloid cells can be differentiated using microglia specific factors and endogenously, this is the natural origin of the resident tissue macrophages of adjacent tissues like the eye or choroid plexus (Gomez Perdiguero et al., 2015; “Principles of resident tissue macrophages revealed by the eye,” 2026). Stem cell derived microglia like cells have been added to a variety of organoid models at a range of time points with varying similarity to human biology.

We generated hiSC derived microglia like cells and assembled them with a neural forebrain organoid model at a time point that recapitulates their endogenous invasion phase. As immune-forebrain assembloids matured, the presence of microglia influenced the maturation of astrocytes, synaptic maturation, tissue level metabolic programs and the overall cellular profile of the copatterned neural tissues. These results highlight a larger neurodevelopmental role for microglia that previously known.

## METHODS

### Microglia differentiation

Microglia differentiation starts from differentiating stem cells to hematopoietic cells, microglial precursors, through the most updated Blurton-Jones protocol(Abud et al., 2017; Hasselmann and Blurton-Jones, 2020; McQuade et al., 2018), commercialized by Stem Cell Technology. Hematopoietic cells have been differentiated in a 12-day process. Briefly, Stem cells were cultured on Matrigel-coated 6-well plates. When cells get to 80-90% confluency and look healthy, dissociation is performed with enzyme-free ReLeSR reagent. Aggregates break down to 100-200 μm in diameter with gentle pipetting, and resuspend 40-80 aggregates per well in a 12-well plate. Cells have been fed with hematopoietic basal media and medium A for the first 2 days, and then medium A was replaced by medium B; half of the media was changed every other day until day 12. On day 12, collect the cells by vigorously pipetting and transferring aggregates into the collection tube, centrifuge, and obtain hematopoietic cell pellets. A flow cytometry assay for CD43, CD45, and CD43 has been performed to confirm the differentiation.

After collecting hpc suspension, 1*10^5-2*10^5 cells per well were cultured in matrigel-coated 6-well plate with 2 ml of microglia differentiation media, and 1 ml was added every other day. On day 12, the suspension was collected and centrifuged to separate cell pellets, then pellets were resuspended in 2 ml of differentiation media, and 1 ml of media was added every other day until day 24. On day 24, differentiated microglia suspension was collected, centrifuged, and cell pellets were resuspended in microglia maturation media for 4-10 days. To keep microglia for more than 10 days, it is necessary to change the media to differentiation media; otherwise, it causes cell death. After the maturation step, microglia differentiation has been verified with CD45, CD11b, and TREM2. Validation by flow cytometry demonstrated essential markers of HPC and microglia differentiation, with 10-20% co-expression of CD34 and CD45, and above 90% CD43 for HPC and 70% co-expression of CD45 and CD11b for microglia

### Brain organoid differentiation

Human cortical organoids were differentiated using a regionalized forebrain differentiation protocol developed by Kadoshima(Kadoshima et al., 2013). H1 cells were dissociated into single cells and then, the cells were plated in GMEM-based neural induction media at a density of 10,000 cells per well in 96-well V-bottom ultra-low adhesion plates. After 18 days, the organoids were transferred to 6-well low-adhesion plates on an orbital shaker at 90 RPM and cultured in DMEM/F12-based media. From day 70, 1X B27 supplement was added to the media. Organoids were fed every other day throughout the culture period(Andrews et al., 2023).

### Assembloid culture

Week 5 brain organoids were transferred to a V-shape 96-well plate on day 0, and fed with 1:1 microglia differentiation media and media 3 of brain organoid, named as assembloid media. On day 1, differentiated microglia have been harvested, and microglia are counted with a cell counter (device name? Countess?), microglia are centrifuged for 10 minutes at 350g, and resuspended in 100ml of assembloid media, and around 100k per organoid is added in each well and incubated for 72 hours. On day 4 Transfer assembloid to a 6-well low attachment plate, place it on the shaker at 90 rpm.

### Harvesting microglia assembloids

Put the 4% PFA out to thaw to fix microglia assembloid and brain organoids for 45 mins on the rocker. Wash the samples with 1xPBS for 30 mins, three times on the rocker at room temperature. Add 1 ml 30% Sucrose to each sample and keep it overnight at 4 °C until the samples sink.

### Embedding

When samples are saturated with 30% OCT, use OCT:30% sucrose solution with a 50:50 ratio for embedding microglia assembloids and brain organoids. Wash samples with the embedding solution and then add the sample to the labeled cryomold. Without making any bubbles, add 1 ml embedding solution and put the mold on dry ice for 10-15 minutes. After the mold embeds fully, transfer it to -80 for storage.

### Immunohistochemistry

First, do cryosectioning, 20 μm thickness, and use charged slides. Bake the slides at 50 °C for 10 mins, Block slices with goat serum at room temperature for an hour, and add primary antibody (antibody names?) and incubate overnight at 4 °C. On the second dat take the slides out and wash 3x with T-PBS (1xPBS + 0.1% Triton) at room temperature. Dilute secondary antibodies and incubate at room temperature for 2 hours, followed by 10 min counterstain with DAPI, then quick wash with 1xPBS-T and wash 3 times with room temperature 1xPBS. At the end, coat with mounting media and cover the slides with a glass coverslip, and after drying (approximately 24 hours) slides are ready to image.

### Water and Lipid Extraction

Crush organoid with dounce or, add 400uL of ice cold methanol and vortex for 30 seconds, then add 200uL of Chloroform and 400uL of DI H2O and vortex for 30 seconds. Sonicate samples for about 5 minutes to dissolve completely. Centrifuge at 16000g for 15 minutes at 4 °C. Extract the top aqueous layer and pipette it into a new tube. Extract the bottom lipid layer in another tube. Put tinfoil over the top of each tube and punch a hole in it with a small pipette tip. Speedvac overnight.

### Mass spectrometry and analysis

All LC-MS experiments were performed on a Thermo Vanquish UPLC-Exploris 240 Orbitrap MS instrument (Waltham, MA). Chromatographic separations were performed in hydrophilic interaction chromatography (HILIC) mode on a Waters XBridge BEH Amide column (150 x 2.1 mm, 2.5 µm particle size, Waters Corporation, Milford, MA). The mobile phase was composed of Solvents A (10 mM ammonium acetate, 10 mM ammonium hydroxide in 95% H_2_O/5% ACN) and B (10 mM ammonium acetate, 10 mM ammonium hydroxide in 95% ACN/5% H_2_O). After the initial 1 min isocratic elution of 90% B, the percentage of Solvent B decreased to 40% at t=11 min. The composition of Solvent B maintained at 40% for 4 min (t=15 min), and then the percentage of B gradually went back to 90%, to prepare for the next injection. Using mass spectrometer equipped with an electrospray ionization (ESI) source, we will collect untargeted data from 70 to 400 m/z. After data collection, the mass isotopomer distributions (MIDs) for each sample were calculated by integrating individual metabolites, and the IsoCor software was employed for the correction of natural abundance of isotopes.

## RESULTS

### Microglia accelerate glial development or proliferation

To assay the broad tissue effects of the addition of microglia we used markers of microglia (Iba1) and a broad marker of glial lineage cells, including radial glia and astrocytes (GFAP). GFAP+ cells are not typically detected in forebrain organoids using this differentiation protocol, but our coculture media mixed microglia and neural factors (Andrews et al., 2023). Iba1+ ESC derived microglia showed robust engraftment and invasion into neural tissues (**Fig 2A**). Glial lineage cells are not reported so early in forebrain organoid development, but were visible proximal to microglia, especially at the tissue edges (**Fig 2B**). Neither microglia nor glial lineage cells could be found in week 6 organoids (**Fig 2C**). By week 5+3 microglia form a Iba1 high population around the periphery with much lower expression within the assembloid (**Fig 2D**). Glial lineage cells remained, but were only detected proximal, or in contact to a microglia (**Fig 2E**). At this later stage, some glial markers expressed in organoids in cells with extended morphologies (**Fig 2F**). Overall the presence of microglia accelerated the maturity of glial cells, but only within cell-cell contact range.

**Figure 1:**
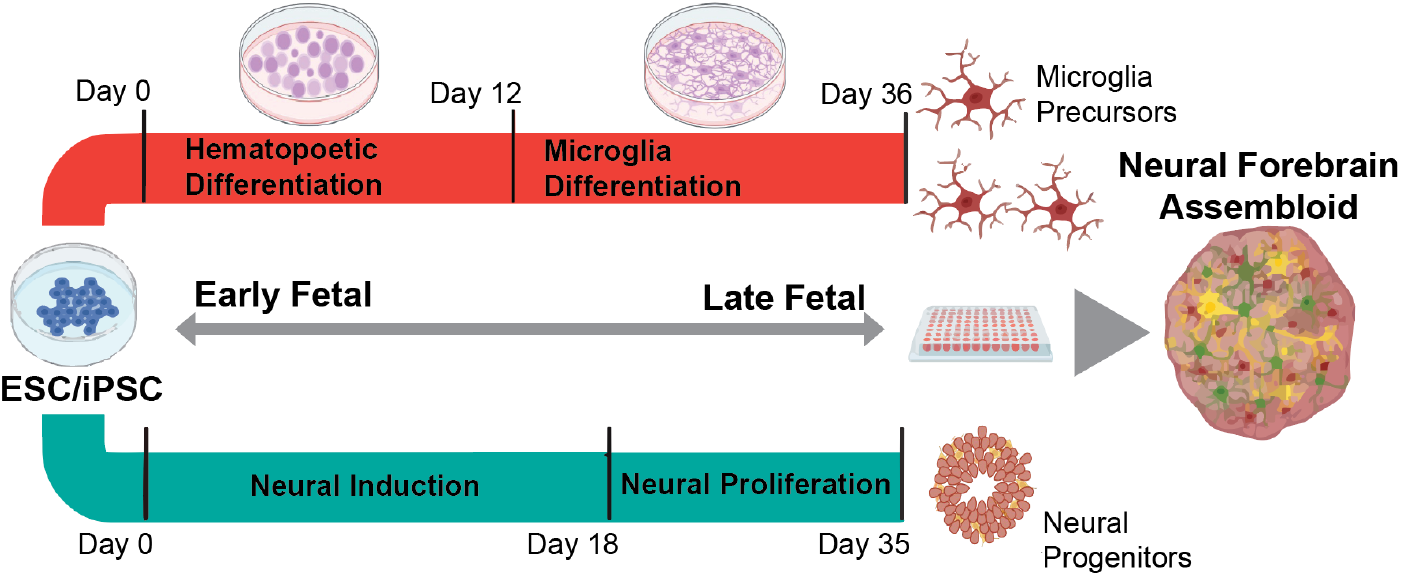
Schematic of parallel culture of neural and microglia precursors followed by assembly. Microglia and neural forebrain tissues were differentiated in parallel, then assembled after neural proliferation, but before maturation and differentiation to post-mitotic neuronal and glial cell types.

**Figure 2:**
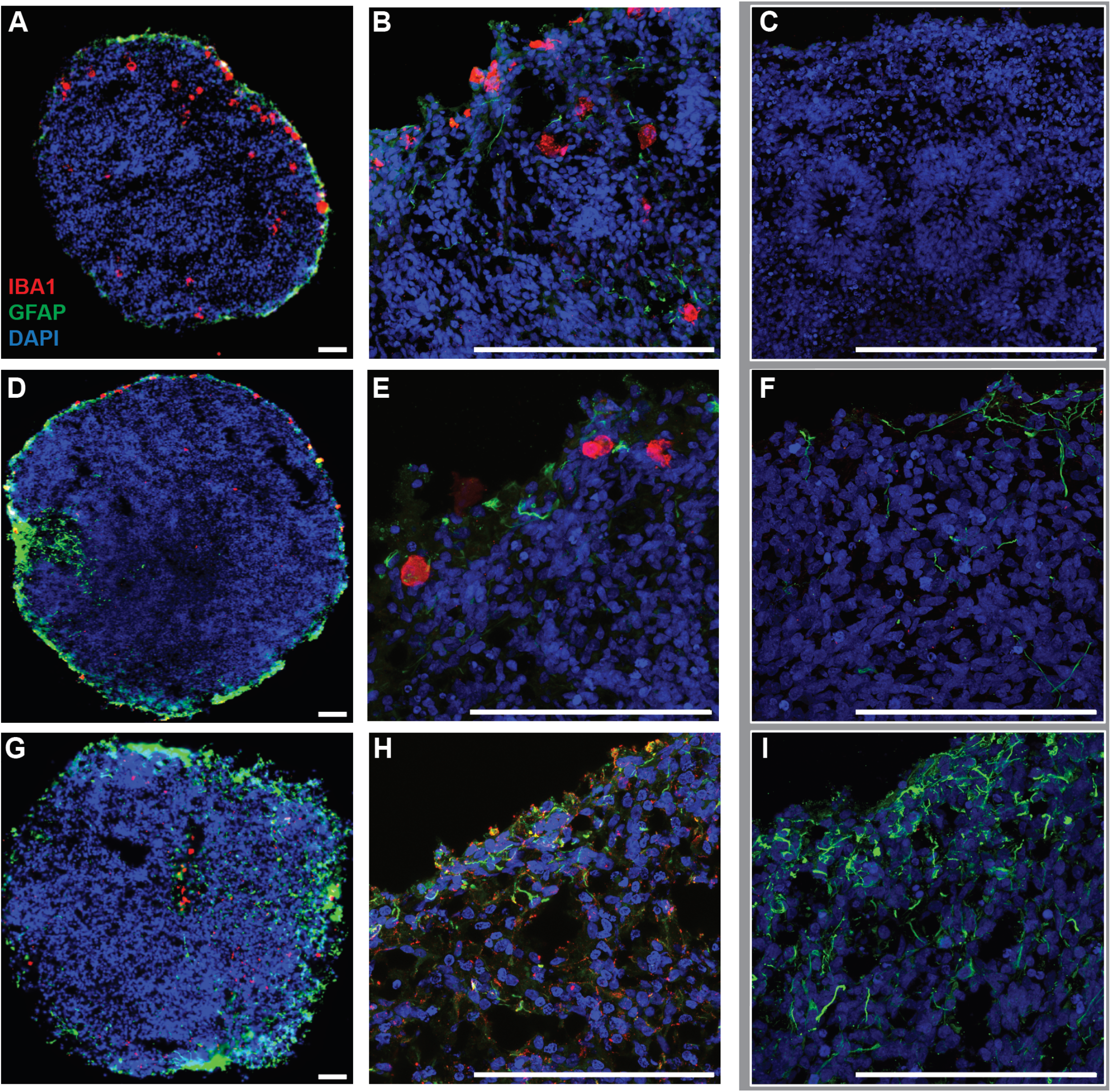
Immunohistology for microglia and glia in week 6-8 forebrain assembloids vs organoids. **(A)** Whole assembloid section showing microglia distribution. **(B)** GFAP+ cells are mostly at the assembloid periphery and proximal to Iba1+ **(C)** Week 5+1 organoids are Iba1- and GFAP-. **(D)** Week 5+3 assembloids are larger and more cell dense. **(E)** GFAP+ cells remain at the assembloid periphery and proximal to Iba1+ **(F)** Week 3 organoids show some GFAP+ cells with no peripheral distribution. **(G)** Week 12 assembloids have abundant Iba1+, GFAP+, NeuN+ cells. **(H)** Near the core of the assembloids Iba1+ and GFAP+ cells are less but remain a common type **(I)** Iba1-organoids have more abundant GFAP+ cells at their periphery.(Scale bar = 25µm)

Looking at a later time point, microglia continued to proliferate in assembloids, appearing deep into the neural tissue with similarly widespread glia, while neuronal lineages became common at the periphery **(Fig 2G)**. Towards the center of the assembloids glia were less common, but present around microglia **(Fig2H)**. In Iba1-organoids, glia became a more common cell type over time, but with few detected neurons **(Fig2I)**.

### Microglia influence glia towards astrocytic fates

Given the early appearance of GFAP+ glia in the microglia-forebrain assembloid, we asked if these cells were differentiating along glial lineages. Using marker for radial glia SOX2, we found this cell fate at the assembloid periphery along with the majority of Iba1+ cells (**Fig 3A**), partially matching the GFAP staining at the same time point (**Fig 2D,E**). Deeper into the tissue SOX2 was less common with cells expressing S100β, a maturity marker for astrocytes (**Fig 3B**). Microglia were uncommon this far into tissues, but were found proximal to or in contact with astrocytes, especially at later time points (**Fig 3 D,E**). In organoids both markers were rare, but could be found by week 3, often overlapping in the same cell type (**Fig 3C,F**).

**Figure 3:**
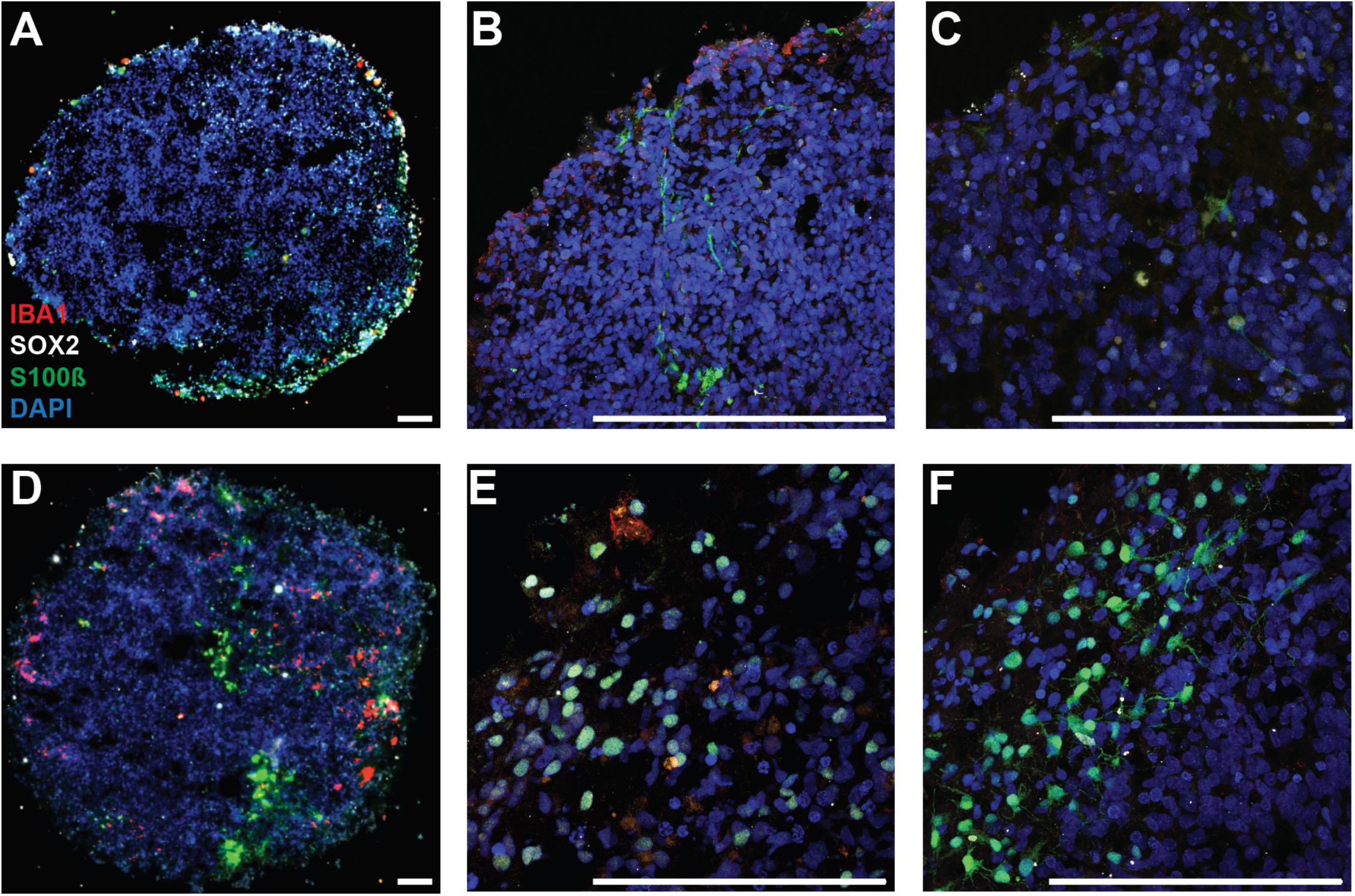
Immunohistology for microglia, astrocytes and radial glia in week 8 forebrain assembloids vs organoids. **(A)**Week 5+3 assembloid distribution of Iba1, SOX2 and S100ß. **(B)** S100ß morphology and proximity to Iba1+ cells. **(C)** Week 5+3 organoids S100β+ and SOX2+ cells are rare. **(D)** SOX2+ cells are mostly gone while S100β+ cells appear deeper into the tissue. **(E)** S100β+ form in clusters proximal to Iba1+ cells. **(F)** Week 5+7 Iba1-organoids now have S100β+ cells. (Scale bar = 25µm)

### Microglia assembloids have enhanced taurine and amino acid metabolism compared to organoids

Neural development includes metabolic transitions as cells become post mitotic (Andrews et al., 2020). Given the enrichment of differentiation and maturity markers in assembloids, we asked how this affected tissue level metabolism. Comparing across all time points gave significant enrichment of metabolites in assembloids and organoids **(Fig 5A)** Metabolite enrichment analysis returned taurine and hypotaurine metabolism as a highly enriched ontology, along with most other amino acids, nicotinamide and nitrogen metabolism in general **(Fig 5B)**.

**Figure 5:**
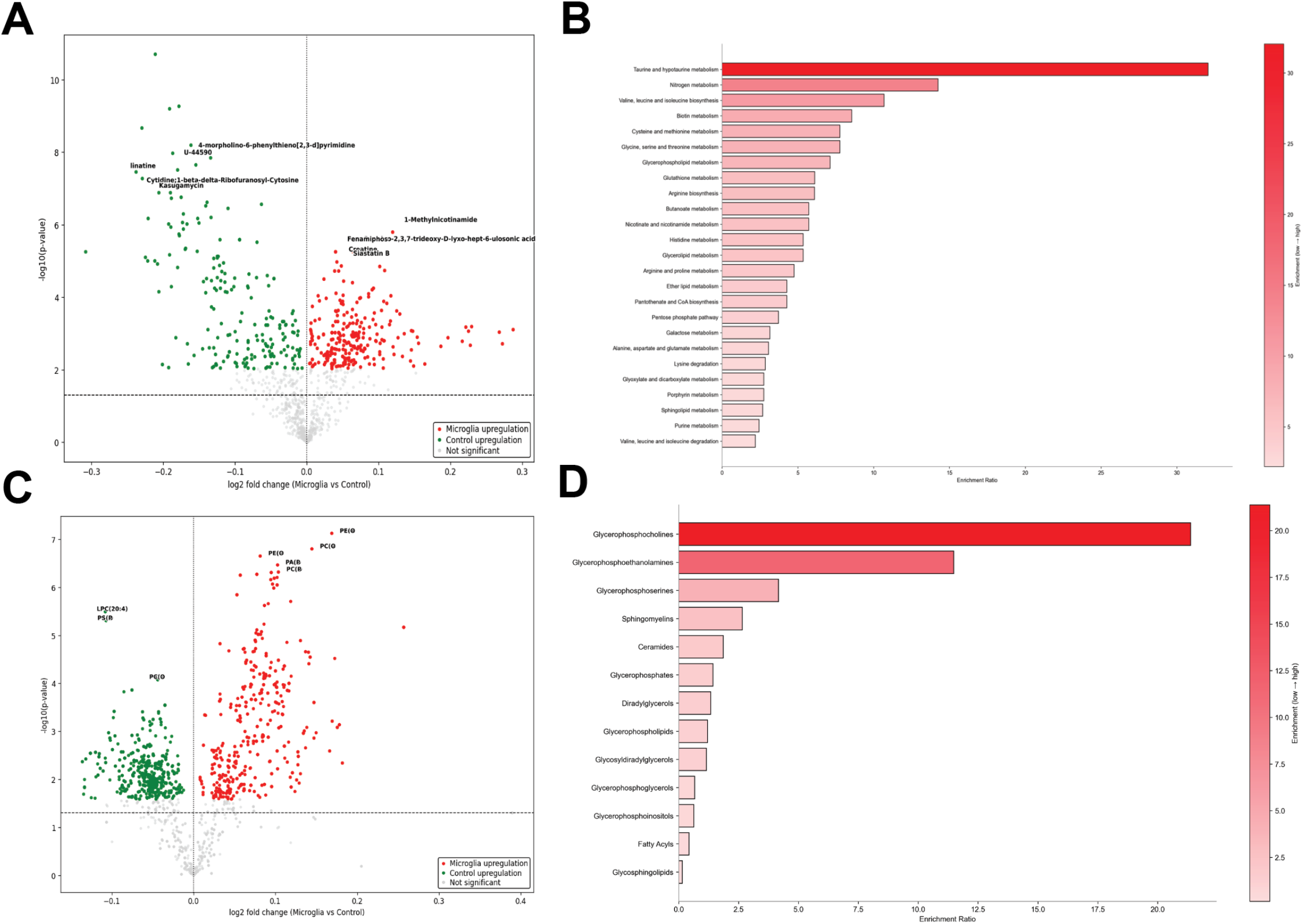
Metabolomic profiling of soluble and lipid metabolites. **(A)** Volcano plot of soluble metabolite abundance between assembloids and organoids. **(B)** Ontological enrichment of differentially abundant soluble metabolites. **(C)** Volcano plot of lipid abundance between assembloids and organoids. **(D)** Ontological enrichment of differentially abundant lipids.

### Microglia assembloids have increased glycerophospholipids, sphingomyelins, and ceramides

A key role of microglia in development and homeostasis is phagocytosis and synaptic pruning, while the primary markers of neuroinflammatory pathology are lipid metabolism genes. Conversely, astrocytes are the primary source of cholesterol in the brain. Given the enrichment of these cell types in the assembloid, we hypothesized that their presence would influence tissue level lipid metabolism. Comparing across all time points, assembloids were highly enriched for multiple lipid species **(Fig 5C)**. Of these, positively charged head groups, glycerophosphocholines and glycerophosphoethanolamines, were the most significantly enriched along with the inner membrane phosphoserine head group. Ceramides were the most microglia associated and expected group to be enriched, but no positively charged ceramides were found, suggesting these lipids came from different cells.

## DISCUSSION

We have built on the growing body of work to generate neuroimmune organoids, work that began with neural forebrain differentiation (Kadoshima et al., 2013), then incorporated primary microglia (Popova et al., 2021), and finally ESC derived microglia-like cells. These advances rightfully questioned whether the newly incorporated microglia developed their endogenous functions (Bennett et al., 2021; Schafer et al., 2023). The human microglia forebrain assembloid supports stable microglial engraftment, migration into inner regions, and accelerates astrocyte maturation in a spatially organized manner; key aspects of human brain development. This work presents some evidence of ESC derived microglia taking on a developmental role, largely influencing the maturation of astrocytes. Evidence of maturity extends to metabolism, where the assembloid tissues show evidence of earlier development of astrocyte functional roles in synapse development.

Microglia-forebrain assembloids accelerate the appearance of GFAP+ cell types to as little as 1 week of co-culture, with markers of maturity appearing by 3 weeks of co-cuture (**Fig 2A,B Fig 3 A,B**). While the reservoir of SOX2+ progenitors does not change, existing glial cells take on an earlier lineage commitment to astrocytes, particularly away from the organoid periphery, an area usually enriched in radial glia cells. This suggests that microglia directly interact with developing glial cells, supported by their high TGF-beta expression.

Metabolomic profiles serve as a powerful molecular assay of function at the tissue level. The presence of microglia altered organoid metabolism beyond the contribution of the added cells alone. Enriched taurine metabolism was an unexpected result with little relation to microglia. Instead, taurine and hypotaurine are produced entirely by astrocytes and function as a developmental neuro/gliatransmitter (Ramírez-Guerrero et al., 2022; Wu and Prentice, 2010). Synthesis of taurine in the central nervous system (CNS) occurs predominantly in astrocytes. A metabolic coupling between astrocytes and neurons has been reported, in which astrocytes provide neurons with hypotaurine as a substrate for taurine production. Taurine has antioxidative, osmoregulatory, and anti-inflammatory functions, among other neuroprotective properties. Mature astrocytes release taurine as a gliotransmitter, both in development and across all stages of life, promoting both extracellular and intracellular effects in neurons.

Altered lipid metabolism was the most expected result, with enrichment of ceramides in the assembloids confirming their production as signaling molecules (McInnis et al., 2024). One result that requires additional studies is the enhancement of positively charged lipid species like glycerophosphocholines and ethanolamines (Chausse et al., 2021). We did not detect enrichment of positively charged head groups in the ceramides, suggesting these lipids are coming from a non-microglia cell type. Deconvolving the lipid profiles of complex tissues is an ongoing challenge for the field (Prakash et al., 2025).

Molecular dysregulation during corticogenesis impacts vulnerability to neurodevelopmental disorders and psychiatric diseases (Bennett et al., 2021). An important, yet underexplored aspect of neurogenesis is the metabolic regulation of cellular development and tissue formation, as alterations in metabolic programs are implicated in a range of neurodevelopmental disorders. In this study of the microglia forebrain assembloid we have found evidence of microglia contributing to the metabolic space that shapes development (Afridi et al., 2020).

## Notes

Declare any conflicts of interest: The authors have no competing financial interests.

Data sharing policy: The raw that support the findings of this study are available from the corresponding author.

### Competing Interest Statement

The authors have declared no competing interest.

